# Generation of Neural Stem/Progenitor-like Cells from Cultured Human Peripheral Blood Mononuclear Cells combined with Cannabidiol

**DOI:** 10.1101/2025.07.18.665629

**Authors:** Saw Kalayar Aye, Sineenat Sripattanakul, Krai Daowtak, Chalermchai Pilapong

## Abstract

Recent advances in chemical reprogramming with small molecule combination have enabled the direct conversion of somatic cells into different cell lineages without genetic modification. This study aimed to investigate reprogramming and differentiation capacity of peripheral blood mononuclear cells (PBMCs) after treating with only a single small molecule, cannabidiol (CBD). The differentiated reprogrammed cells exhibited high expression of neuronal stem/progenitor cell (NSPCs) markers without pluripotency markers, suggesting cellular identity was switched to NSPCs via direct reprogramming. Transcriptomics and proteomic analyses of the differentiated reprogrammed cells showed the remarkable expression of genes specific to NSPCs and endocannabinoid system along with regenerative parameters and morphogenesis. Unexpectedly, we have found that PBMCs may inherently possess certain levels of plasticity or potency, possibly through dedifferentiation or transdifferentiation mechanisms. Our findings convey the idea that CBD together with intrinsic plasticity of PBMCs might be able to induce the transdifferentiation of PBMCs.

## Introduction

Degenerative and aging diseases, resulting from continuous loss of functioning cells along with poor regenerative capacity, contribute to high morbidity/mortality rates due to no successful curative therapies. Since adult mammals have scanty residing stem cells, the induction of dedifferentiation and transdifferentiation is important to obtain cellular sources of regeneration (*1, 2*). Cell reprogramming, a ground breaking technique introduced by Yamanaka (*3*) over a decade ago, is another approach to get enormous number of stem cells and they are named as induced pluripotent stem cells (iPSCs). All these three routes (dedifferentiation, transdifferentiation and reprogramming) lead to the establishment of powerful regenerative medicine (*4-7*). Regenerative medicine can replace lost cells by providing stem cells or their differentiated derivatives, which can be produced either through the proliferation of preexisting endogenous stem/precursor cells or by the injection of exogenous stem/progenitor cells (*8*).

Peripheral blood contains certain kinds of stem cells including mesenchymal stem cells (MSCs), hematopoietic stem cells (HSCs), endothelial progenitor cells (EPCs), very small embryonic like stem cells (VSELs), etc. (*9*). Scientists have been highly enthusiastic to isolate and expand these stem cell populations of human peripheral blood, which are in a very small amount in normal equilibrium situations, aiming for autologous regenerative medicine (*9, 10*). Moreover, the immune cell within peripheral blood possesses a regenerative potential by modulating stem cell niches through immunity (*8, 11*). For cells to survive, develop, and function properly, an inflammatory microenvironment mediated by the immune response is necessary (*12*). A microenvironment that is conducive to regeneration requires appropriate immune responses (*11, 12*). Human peripheral blood cells can provide significant in vitro cell multiplication and can serve as better donor cells for cell reprogramming. The non-invasive accessibility of peripheral blood from patients also makes it highly attractive for personalized regenerative medicine. Studies have been proved that nearly every cell type derived from the three embryonic layers, including blood cells, endothelial cells (ECs), hepatocytes, cardiomyogenic cells, muscle cells, epithelial cells, and neural cells can be differentiated from PBMCs (*13, 14*). In addition, the ability of PBMCs to become iPSCs increases their capacity for phenotypic conversion. With the advancement of techniques for isolating and purifying specific PBMC subpopulations, it is now possible to get pure populations from the PBMC fraction, and the research works showed that even one subpopulation of PBMC can give rise to other somatic cells by cell reprogramming. For example, transdiffererntiation of neuron like cells from monocytes (*15*), and from T cells (*16*) were reported since 5 years ago. The stem cell field is growing rapidly and newer avenues for reprogramming related regenerative mechanisms by using PBMCs have been continuously developing.

Cannabinoids (CBs) and endocannabinoid system (ECS) have gained much attention due to their potential therapeutic benefits, including for degenerative disorders (*17-20*). The ECS, all together with other components of it such as endocannabinoids, synthesizing and degrading enzymes, receptors, and transport proteins, which can be found in almost all organisms including unicellular eukaryotes to humans, mediate the activity of cellular functions, and homeostasis (*19*). ECS levels can determine developmental processes from implantation to organs formation, especially brain (*17, 21-23*). The contribution of ECS will continue even after birth and it remains involved in adult regeneration processes (bone turnover, neural regeneration, wound healing, etc.). The ECS can influence regenerative parameters like cell migration, proliferation, survival, and inflammation (*24*). CBD, one kind of phytocannabinoid (pCB), with positive effects on chronic major organ disease, aging, epilepsy, pain disorder, to sleep deprivation, can also regulate ECS via both cannabinoid receptors and other ECS pathways (*25-27*). CBD can enhance regenerative capacity similarly to ECS and these effects are in dose-dependent manners according to recent studies (*24, 28*). On PBMCs, CBD can modulate immune reactions and give mainly immunosuppressive effects (*29-31*), providing regenerative permissive microenvironment for tissue remodeling. This immunomodulating effect together with the high regenerative potency of CBD raises CBD to be a great potential for newer research in regenerative field. Particularly, for reprogramming approach, the functions and roles of CBD in this context are still in its infancy, and exploring this capability of CBD on somatic cells through ECS can open the next chapter of CBD and ECS in regenerative medicine.

Above mention substantiates the potentiating effects of CBD and ECS on regenerative parameters of stem cells. PBMC itself also possesses high regenerative properties by giving multi-differentiation capacity and niche modulating action for regenerative permissive microenvironment. Treating ECS rich PBMCs with CBD might become a promising approach for regenerative medicine and attractive candidates for further research to introduce a novel reprogramming approach.

## Results and discussion

### CBD triggers reprogramming and differentiation potency of cultured PBMCs into neural stem/progenitor cells (NSPCs) by priming transdifferentiation

As a great discovery and application of ECS and CBD in the field of regenerative medicine, our main focus was to find out whether CBD was capable of driving cell reprogramming and differentiation of human PBMC. At Frist, 1 million PBMCs were typically seeded in the 6-well plates containing RPMI1640 media comprising CBD, and cell differentiation pattern was monitored over time. After 7 days of culture in the presence of CBD, a significantly greater number of colonies and differentiated cells were observed compared to the untreated control (Figure 1A), indicating the induction possibility of CBD towards reprogramming and differentiation of PBMCs. Additional microscope images of untreated cells and CBD treated cells for 7 days from different donors are shown in figure S1. As a number of colonies were observed in CBD-treated cells, and colony formation can be considered as one of the characteristics of stem cells and neurospheres. Subsequently, we decided to check different markers of pluripotency stem cells (PSCs) (*32, 33*) and neural stem/progenitor cells (NSPCs) (*34, 35*) by means of alkaline phosphatase assay and immunofluorescent staining. As the results show in Figure 1B, the colony highly expressed typical markers of NSPCs such as SOX2 and Musashi-1, while there was less expression of Nestin and PAX6. Regarding pluripotency markers, no expression of TRA-1-81, OCT4, and alkaline phosphatase was observed. The ability to express NSPC markers but not pluripotency markers indicates that the cells could possibly be directly reprogramed to ectoderm lineages without going through a pluripotency state. To further confirm this observation, transcriptomic analysis was performed. Differential gene expression (DEG) analysis revealed that cells treated with CBD for 7 days showed significant upregulation of genes associated with fetal NSPCs (Figure 2A, left panel and table S1). Protein-protein interaction (PPI) analysis of the upregulated NSPC DEGs showed that genes at regulatory network nodes play crucial biology and function of NSPCs (e.g. *NES, FGF2, FOS, TP53* for NPCs differentiation and nervous system development(*36*); *NOTCH1* for maintenance of NSC (*37, 38*); *PAX6* for neuroectoderm and neural rosette cells fate determinant (*38, 39*); *TUBB* for development, differentiation and maintenance of neurons (*40*);, *CD44* for radial glia fates (*38*);, *Vimentin* for neurodevelopment (*41*); *NCAM1* for synapse formation (*42*), (Figure 2A, top right panel). With an enrichment of GO term, neuroepithelial cell differentiation, glial cell differentiation, gliogenesis, regulation of neural precursor cell proliferation etc. were significantly remarked (Figure 2A, bottom right panel). In addition to fetal NSC/NPC, adult NSC genes (*40, 43*) were also found to upregulate in the cells treated with CBD for 7 days (Figure 2, left panel and table S2). PPI network with GO enrichment also showed that CBD was capable of inducing biological processes related to glial cell development and gliogenesis. At the same time, a number of ECS-related genes were also significantly expressed in the cells treated with CBD, compared to untreated cells (Figure 2C, left panel and table S3). Based on PPI analysis of enriched ECS genes (Figure 2C, middle panel), CBD can activate ECS gene network, and core regulatory network genes (nodes) were found to involve in their receptors and transporters, as well as their synthesizing and degrading enzymes. GO enrichment based on enriched ECS genes suggested that CBD can activate lipid catabolism and metabolism, which could be a crucial biological process regulating reprograming and differentiation of PBMC (*44*) (Figure 2C, right panel). As is known, ECS is enormously found in the central nervous system (CNS) and plays an important role in both embryonic and adult neurogenesis by enhancing proliferation, self-renewal, migration, and cell survival of the neural cells at different stages (*21, 22, 45*). Based on these observations, we hypothesized that CBD may activate ECS biology involving in the PBMC reprogramming and differentiation. According to the PPI network of enriched ECS and NSPCs shown in figure S2, coordination between ECS and NSPCs were observed, emphasizing that reprogramming PBMCs into the NSPCs could be go through a mechanism effectuated by endocannabinoid system inherited within the PBMCs. Notably, there was direct coordination between Tumor Protein P53 gene (*TP53*) and peroxisome proliferator-activated receptor gamma gene (*PPARG*), belonging to NSPC and ECS, respectively. *TP53* was previously found to promote lineage commitment of human embryonic stem cells (*46, 47*), while *PPARG* was found to play an important role in lipid metabolism (*48*) and in regulating different biology and function of NSPCs (*49-52*). Potentially, these two genes could be one of the key players mediating reprogramming and differentiation of PBMCs induced by CBD. However, the mechanistic explanation on this aspect is still unclear, especially the link between the endocannabinoid system and the hematopoietic system.

**Figure 1.**
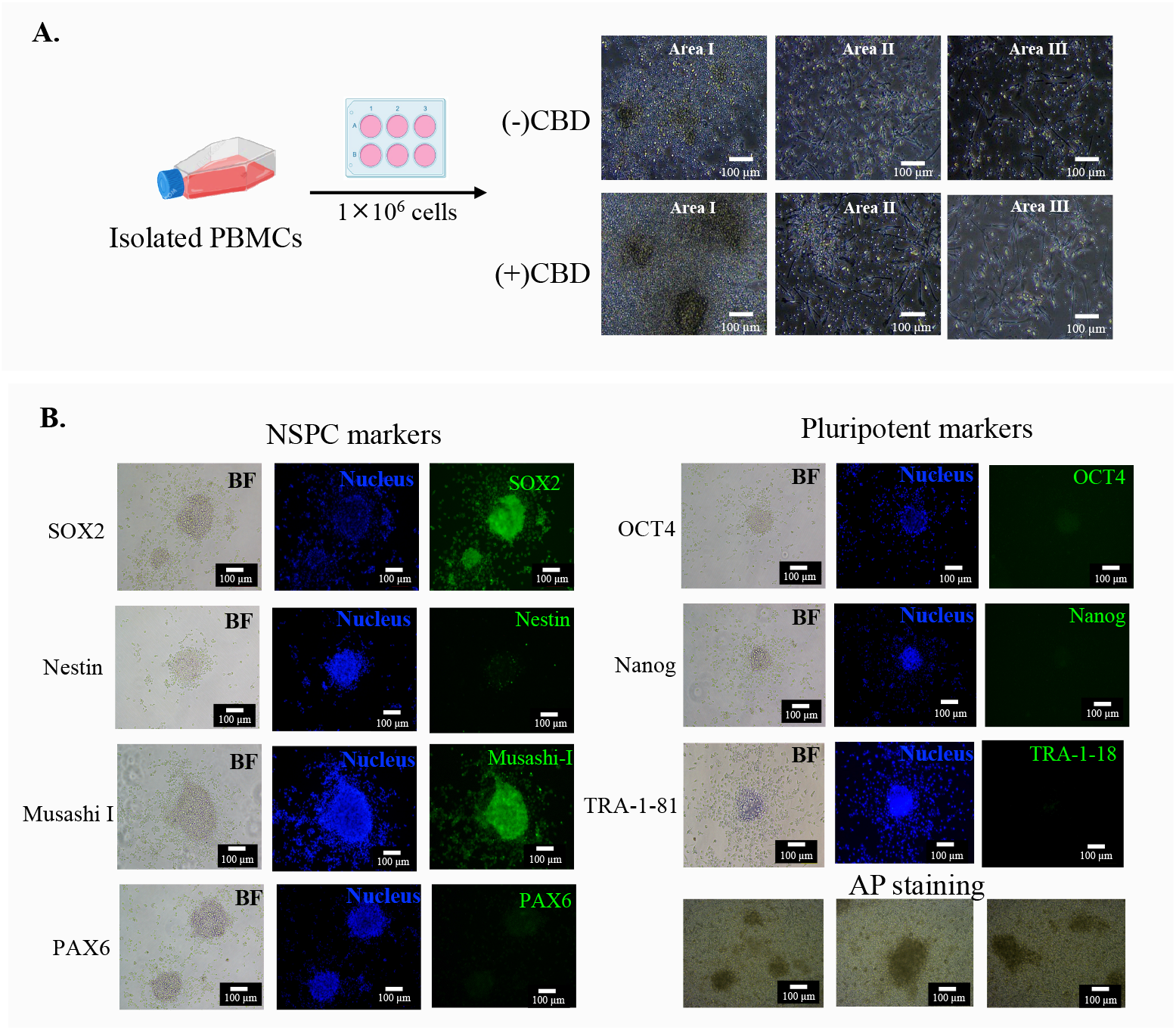
Cultured PBMCs revealed positive markers of NSPC. (**A**). Schematic illustration of cell culture protocol of PBMCs and typical microscopic images (with different areas) of PBMCs seeded in 6-well plates for 7 days with and without CBD treatment. (**B**). Immunofluorescent staining for NSPC, pluripotent markers, and alkaline phosphatase staining of the cell colonies obtained from CBD treatment.

**Figure 2.**
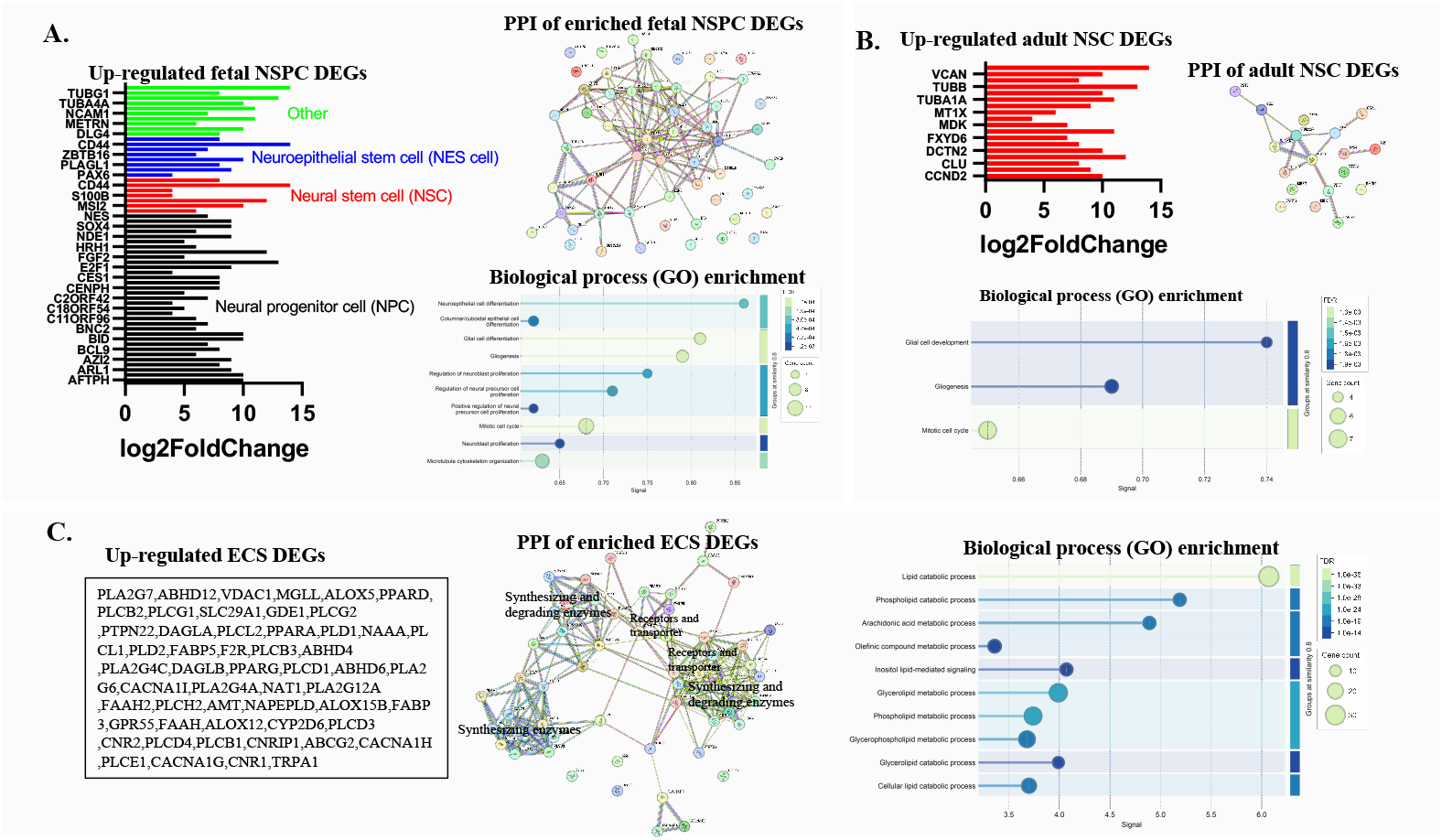
Significant gene expressions of seven days cultured CBD treated PBMCs compared to untreated cells, in 6-well plate. Upregulated DEGs along with the corresponding PPI networks (medium confidence - 0.4) and GO enrichment terms related to (**A**) fetal NSPC and (**B**) adult NSC respectively. (**C**) ECS genes expressed in the CBD treated cells. (p < 0.001).

When further culturing the PBMCs in the presence of CBD for longer time, spindle-shaped cells with different elongation were typically observed (Figure 3A), as the time passed (additional microscope images of untreated cells and CBD treated cells from different donors are shown in figure S3). Typical images of long-spindle shape cells derived from CBD treated PBMCs are shown in Figure 3B (from 3 different individuals). Notably, the length of the cells could be up to submilliter after incubating for over 20 days). Based on the observation regarding an expression of NSPC markers described above, we further checked an expression of different NSPC markers including SOX2, Nestin, PAX6, Musashi-1, Vimentin and GFAP in long-spindle shape cells using IF staining. As the results show in Figure 3C, the differentiated cells show highly stained-biomarkers for PAX6, Vimentin and GFAP, and moderately stained-biomarkers for SOX2, Nestin, Musashi-1 (additional IF results from different donors are shown in figure S4). The presence of such biomarkers in the differentiated cells along with a thin and elongate spindle shape could be an indicator of radial glia (RG) cells, bipolar-shaped neural progenitor cells that are responsible for producing different brain cells including astrocyte, oligodendrocytes, and neuron (*53-57*). We further checked an expression of gliogenic transcription factor nuclear factor I (NF-1) (*58, 59*), and neuronal markers beta-tubulin 3 (TUBB3) (*60-62*) and synapsin (*63, 64*) in a thin, long spindle cells, as expected, negative expression for those markers were observed in the differentiated cells (figure S5), confirming the cells did not undergo astrogliogenesis and neurogenesis.

**Figure 3.**
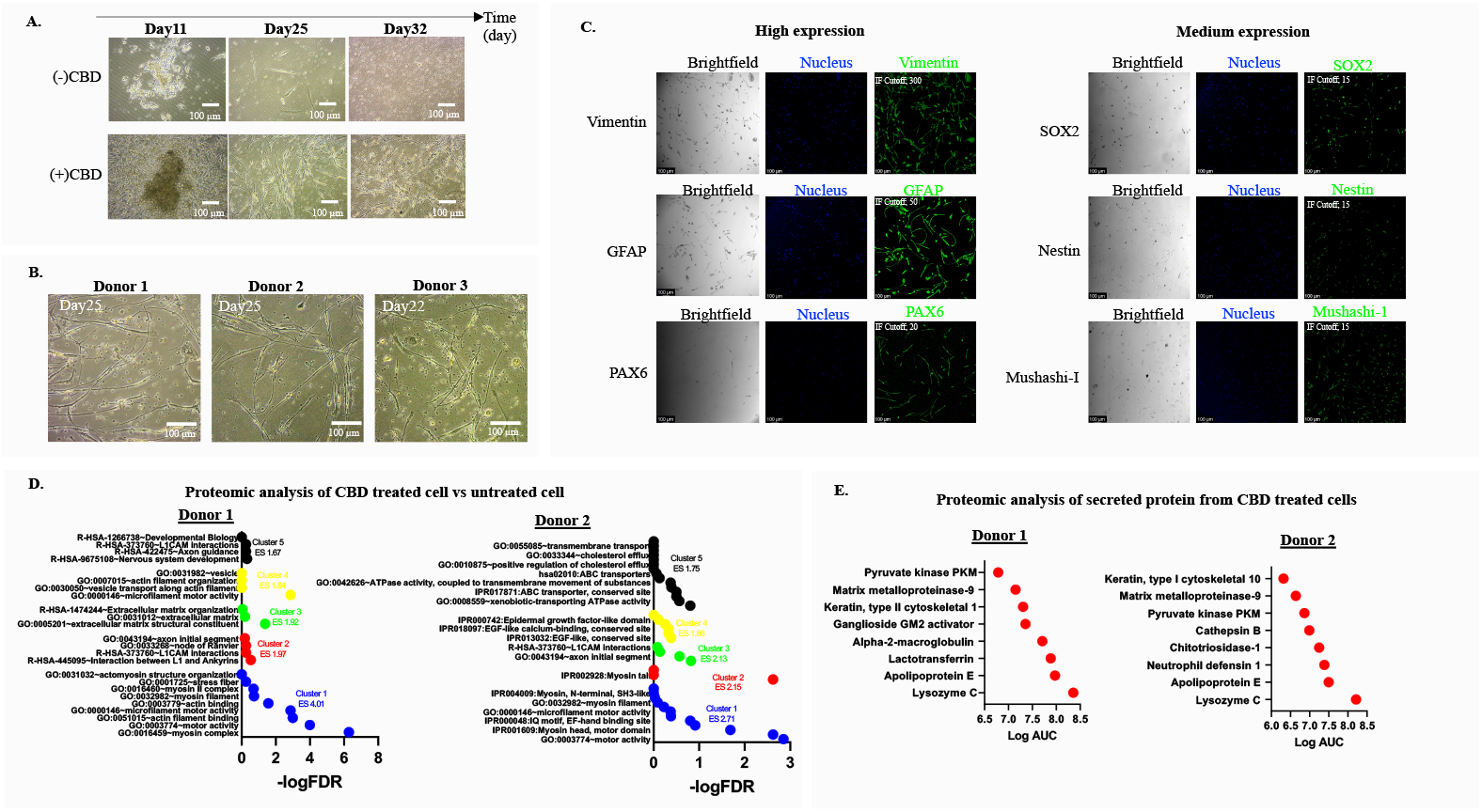
Differentiated cells coming from CBD treated PBMCs express NSPC markers and cytoskeleton based neurodevelopmental related biological processes. (**A**) Typical microscopic images of cells treated with CBD at different lengths of time. (**B**) Microscopic images of differentiated cells obtained from CBD treated PBMCs for more than 20 days from 3 different donors. (**C**) Confocal images of immunofluorescent staining for NSPC markers: Vimentin, GFAP, PAX6, SOX2, Nestin and Musashi-1 in differentiated cells (thin and long spindle shape cells) obtained from CBD treated PBMCs for more than 20 days. (**D**) Different DAVID annotated clusters based on proteomic analysis of the differentiated cells obtained from CBD treated PBMCs for 39 days, compared to cells that have not been treated with CBD (2 independent donors). (**E**) Proteomic analysis of secreted proteins in cell culture media collected from CBD treated cells for 39 days (2 independent donors).

Proteomic analysis of differentiated cells being incubated with CBD with respect to cells that have not been treated with CBD for 39 days of two independent donors revealed that more than 300 proteins were significantly expressed in differentiated cells (table S4). By conducting DAVID annotation, the top annotation cluster was found to associate with biology and function of the cytoskeleton, such as myosin and actin, which was consistent with the characteristics of cells that had the appearance of elongated spindle-shaped cells. In addition, proteins related to epidermal growth factor, node of Ranvier, axon initial segment, interaction between L1 and Ankyrin, L1CAM interaction, axon guidance, nervous system development, and developmental biology, etc. were also found to be upregulated (Figure 3D). This proteomic analysis suggested that the differentiation of PBMCs induced by CBD involved the development of cells with high cytoskeleton activity, including cells with processes related to epidermal growth factor or processes related to the nervous system. At the same time, cell culture media of the differentiated cell being incubated with CBD from two independent donors were also collected for secreted protein identification. As the results shown in Figure 3E, several new proteins were found in the culture media cells being incubated with CBD. Interestingly, apolipoprotein E (APOE) was found to be one of the most abundant secreted proteins found in the media. Although APOE can be expressed at high levels under stress or disease conditions, APOE can be highly expressed in Nestin/GFAP double-positive NSPCs and play a role in NPC maintenance (*65, 66*). Taken together, capable of secreting APOE could support the notion that the elongated spindle-shaped cells could behave like RG cells.

The above-mentioned results showed that culturing the PBMCs in the presence of CBD over a period of time was able to turn the cultured PBMCs into differentiated reprogrammed cells having characteristics of NSPCs. Preferably, the predetermined concentration of CBD was 1 to 5.0 μg/mL, which is in a range of non-toxic dose, based on cell viability assay (figure S6). It should be noted that expression of CBD specific-receptors or transports might be different in individual, inferring that biological response of PBMCs to CBD could also be individual differences. By measuring a typical target of CBD (CB2 receptor), as expected, it was found to be expressed differently in independent donors (figure S7). In fact, the optimum dose of CBD (e.g. obtaining a long spindle shape cells) might not be the same in different donors, but effective dose was in between a dose response range of 1-5 μg/mL. According to our observations of the cells from different donors and sources, almost all the cells behave using the same differentiation patterns. Typical microscopic images of the differentiated cells induced by different doses of CBD between different donors are shown in figure S8.

### Enriched regenerative properties e.g. differentiation, development, stem cell in CBD treated PBMCs

Based on the observation cell colonies during the early days of incubation, a certain number of cells were cultured in a lower-surface-area well (96-well plates) at different incubation times. Notably, cell differentiation was not clearly observed, but the cells had a tendency to aggregate and form loosely clustered structures (Figure 4A, top left panel), which are so-called mononuclear clusters (MCCs). In contrast, the colonies obtained from the cell being cultured in larger-surface-area well (6-well plate) comprised of highly packed cells and attached to the plastic plate (as shown in Figure 4 bottom left panel, also Figure 1A, and figure S1). Generally, MCCs or colonies can be formed by various factors, and the exact mechanism is not yet known. However, their formation can be considered as one of characteristics of stemness and cellular regeneration (*24, 67*). Herein, we did not intend to investigate the mechanisms underlying the formation and differentiation but rather sought to determine whether the formation of MCCs, or colonies, and the cell differentiation induced by CBD, was associated with activation of stem cell biology and regenerative property of PBMCs or not. As the results shown in Figure 4 middle top/bottom panel and table S5, either culturing in lower or larger spaces, enhanced regeneration mechanisms including stem cell self-renewal and multi/pluripotency, and cell proliferation were enriched in cells treated with CBD (at day 1 and day 8 for culturing in 96-well plate, day 7 for culturing in 6-well plate) compared to the untreated cells. Similarly, GO enrichment terms showed that the terms related to cell differentiation, cellular development, and stem cell were markedly enriched (Figure 4 right top/bottom panel, and table S6). Based on the above transcriptomic data, it can be implied that CBD are capable of activating genes associated with regenerative capability, which are supposed to be involved in PBMC reprogramming and differentiation.

**Figure 4.**
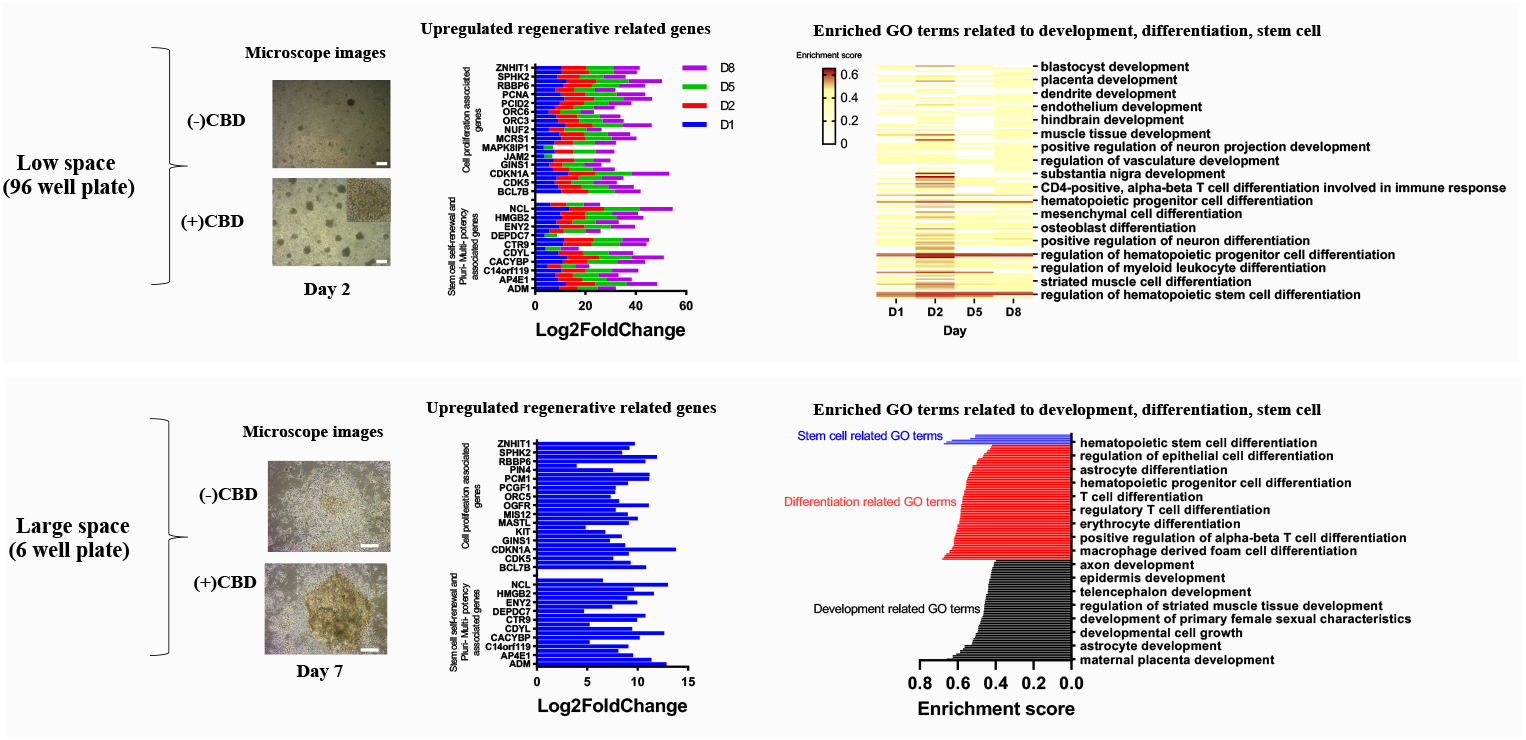
CBD treated PBMCs occupy enriched regenerative properties. Microscopic images of cultured PBMCs together with the expression of upregulated genes in log2 fold change related to the increased regenerative mechanism, (padj < 0.001), and enriched GO terms related to development, differentiation, stem cell (padj < 0.001), of PBMCs treated with CBD in 96-well plates at different time intervals, namely day 1 (D1), day 2 (D2), day 5 (D5) and day 8 (D8) (top panel) and in a 6-well plate at day 7 (bottom panel), compared to untreated cells.

### PBMCs themselves possess the regenerative solid capacity with intrinsic plasticity

From the above results, it was noticed that PBMCs themselves also showed reprogramming and differentiation potencies (but less efficacy by comparing with CBD induction). Once again, we investigated cell differentiation behavior and analyzed stem cell biology and cellular regeneration properties of cultured PBMCs under basic culture conditions without adding CBD. In typical observation, MCC formation was observed when 1 million PBMCs were cultured in lower-surface-area (96-well plate), in a time-dependent manner. Once we cultured the cells in larger-surface-area (6-well plate), colonies can be observed during the early days of incubation. Later, the cells started to differentiate to be spindle-shaped cells (Figure 5A). According to our observations of the cells from different donors and sources, almost all the cells behave using the same differentiation patterns (see additional images from several independent donors are shown in untreated condition of figure S1, and figure S8). By characterizing the colonies and differentiated cells by mean of IF, the colony was still found to express NSPC markers of SOX2, Musashi-1, and PAX6 (but not for pluripotency markers like TRA-1-81, OCT4) (Figure 5B), while the differentiated cells also express RG cell markers such as Vimentin, GFAP, PAX6 and SOX2 (Figure 5C). Above observation reinforces the notion that PBMCs themselves have a property of plasticity toward differentiation. To explore the intrinsic plasticity of PBMCs, RNA seq analysis of PBMC cultured in different cell culture plates (96-well plate and 6-well plate) was carried out. DEGs analysis revealed that various genes orchestrating different lineage identities and pluripotency were expressed in cultured PBMCs (Figure 6A, table S7). By conducting GO enrichment on the transcriptomic data, it was found that a number of terms related to development, differentiation and stem cell have been enriched in cultured PBMCs, as shown in Figure 6B. Similarly, the targeted genes belonging to regeneration capability including stem cell self-renewal and multi/pluripotency, and cell proliferation were also found to have been activated as shown in Figure 6C and table S2. Based on the above transcriptomic data, it was hypothesized that PBMCs might have their own potency and plasticity. It is well known that the majority of PBMCs are immune cells which mostly consist of lymphocytes and monocytes, with a much lesser amount of stem cells, which are so-called PBSCs. Our results show that a number of cells can easily undergo reprogramming and differentiation under basic culture conditions without adding any factors, indicating that PBMCs themselves have a property of plasticity toward differentiation. Based on IF results, we did not observe positive expression of various PSC markers, indicating that transdifferentiation or dedifferentiation could be considered as plasticity mechanisms of PBMCs. However, various genes orchestrating pluripotency found in cultured PBMCs could suggest that the reprograming or differentiation could partially go through pluripotency state somehow. It is important to note that the state of pluripotent, semi-pluripotent or partial pluripotent can be variably attained through different culturing conditions, culturing periods, cell donors, additives and/or reagent administered into the culture medium, the timing that the additives and/or reagent administered into the culture medium, etc. This observation could lead to a new paradigm on PBMCs, suggesting they acts not only as immune cells but also stem-like cells. They could possibly have dual properties existing between immune cells and stem-like cells. Under suitable conditions, PBMCs can activate its own stemness character via dedifferentiation or can prime for differentiation through transdifferentiation. The expression of NSPC represents typical plasticity of the cells. This lineage switching from PBMCs to NSPC could provide additional evidence supporting the role of circulating immune cell for keeping brain homeostasis and plasticity (*68, 69*). Whether this behavior already exists in our bodies is still not known. If so, interestingly, this could be involved in the regeneration mechanisms taking place throughout our whole body because PBMCs can be circulated to all parts of the body. Unfortunately, it is still not fully known exactly what mechanisms are involved and no evidence in living subjects has been demonstrated.

**Figure 5.**
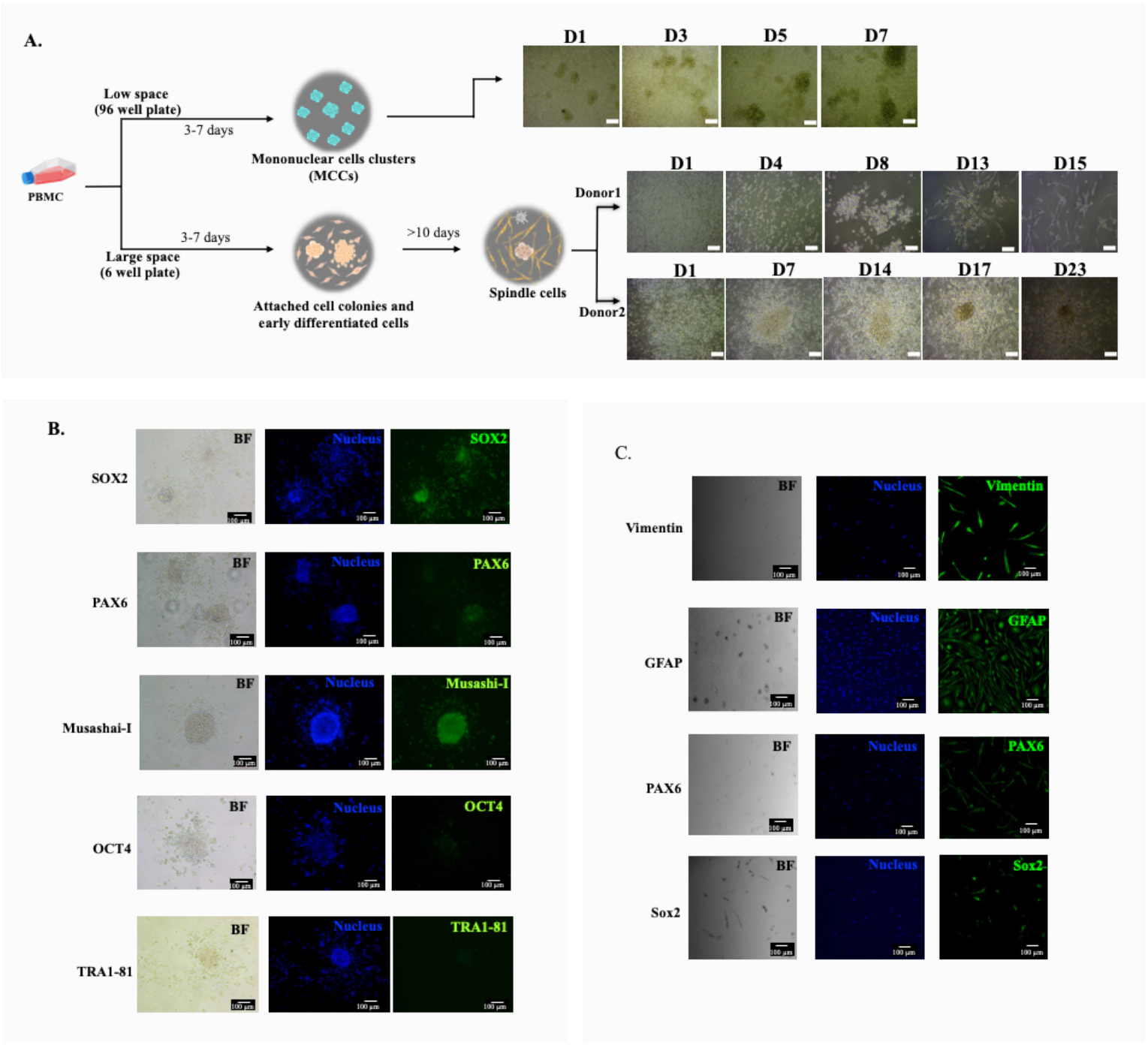
Capability of PBMCs toward differentiation without CBD treatment. (**A**) Schematic illustration of cell culture protocol of PBMCs, typical cell differentiation behavior along with microscopic images of PBMCs seeded in 96-well plates for 1 day (D1), 3 days (D3), 5 days (D5) and 7 days (D7); and in 6-well plates (from two donors) for different lengths of time (scale bar = 100 µm). (**B**) Immunofluorescent staining of the cell colonies. (**C**) Immunofluorescence staining of the differentiated cells.

**Figure 6.**
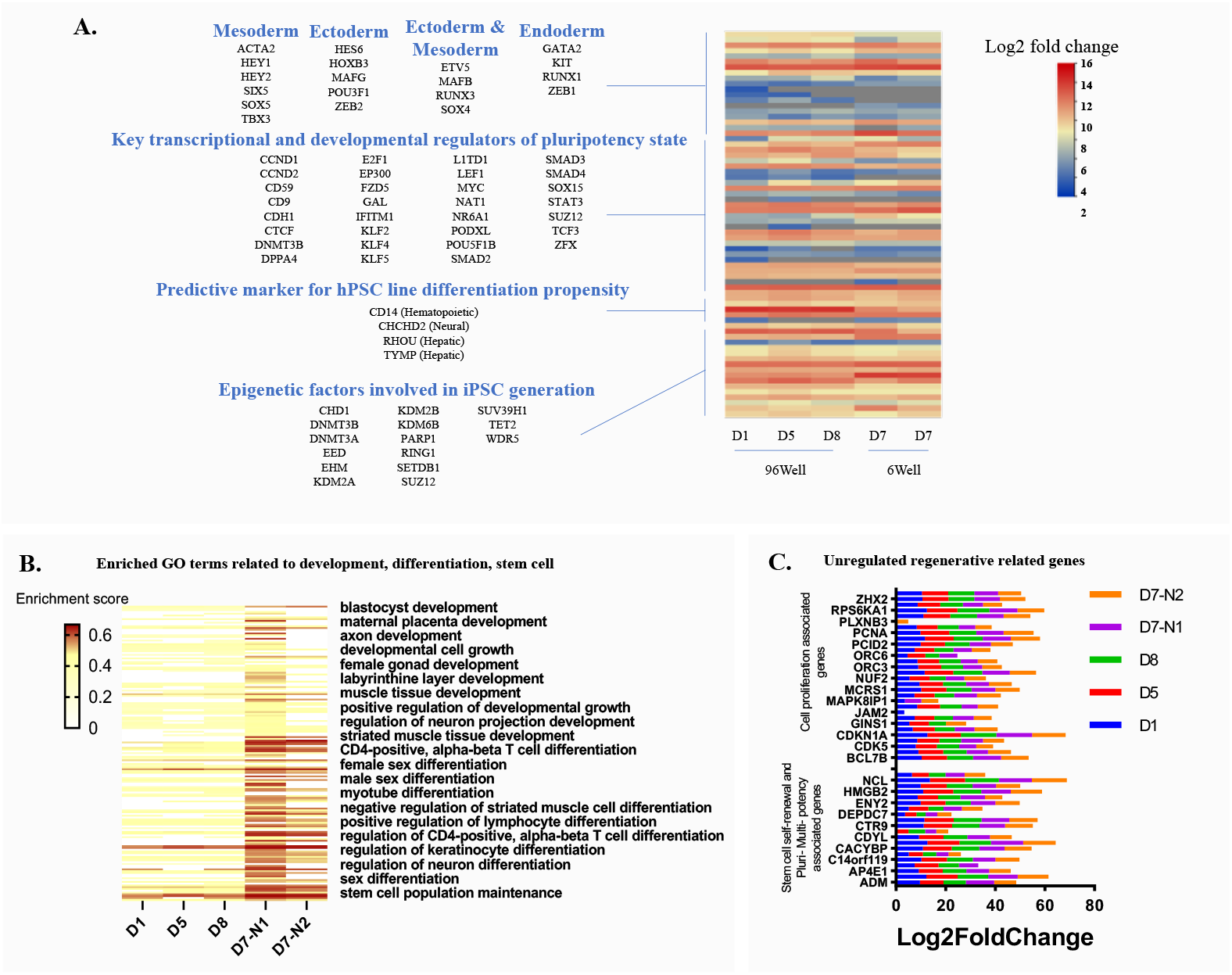
Regenerative solid capacity of PBMCs with their intrinsic plasticity. (**A**) Clustering of lineage –specific gene expression and pluripotency related genes expression (padj < 0.001). (**B**) typical enriched GO terms related to development, differentiation and stem cells (padj < 0.001). and (**C**) Regeneration related DEGs (padj < 0.001) in cultured PBMCs in 96-well plates (D1, D5, D8) and 6-well plates (D7) with two different donors.

### Both PBMC and CBD are critical

Above results showed that PBMCs with and without treatment with CBD were capable of generating NSPC colonies and RGs-like cells. Based on above investigation, CBD itself could not solely play a role on reprogramming and differentiation but intrinsic plasticity/regenerative properties of PBMC also critical because PBMC itself possesses multi-differentiation capacity and niche modulating action for regenerative permissive microenvironment. Similar to previous studies, CBD was used to enhance activity of adipose-derived stem cells (ASCs) and bone marrow-derived stem cells (BMDSCs), focusing on three major regenerative properties: wound healing, migration, and proliferation (*24*). In addition, induction possibility of human gingival mesenchymal stromal cells (hGMSCs) to neural lineage after treating with CBD was also remarked (*28*). In our experiment, the cells treated with CBD were likely to be better at cell reprogramming and differentiation efficiency. Furthermore, the differentiated cells from CBD induction can survive in vitro for several months. The targeted cells obtained from CBD induction has more mutuality and would have more efficient in promoting further differentiation. Whether our NSPC colonies or RG-like cells could produce different neuronal lineages is still ongoing work. Indeed, additional input including biochemical, biophysical, and physiological or biomechanical cues need to be strongly considered for fate immaturity, and to efficiently generate different types of terminally differentiated cells.

## Conclusions

this is the first study to induce direct reprogramming and differentiation of PBMCs by using a single small molecule, CBD. This can provide a novel way of small molecule-based PBMC reprogramming and can be applicable in cell reprogramming technology. This method finally yields target cells of preferred type of NSPCs, which will be advancing cell-based therapy for neurodegenerative diseases. The small molecule that has been used in this study, CBD, can also imitate an endogenous ligand in the human body and have systemic impacts on the body’s homeostatic mechanisms. Therefore, the establishment of this study additionally can serve as a paradigm for the natural intrinsic cellular biological processes and fundamental mechanisms of human beings in the aspect of regeneration. In addition, our results unveiled that PBMCs might have their own plasticity. This study could pave the way to a new paradigm on PBMCs, which may hold potential as a vital mechanism underlying adult regeneration and reprogramming.

**References and Notes**

## Acknowledgments

We would like to thank the volunteer, Blood Bank Unit of Chiang Mai University Hospitals and Naresuan University Hospitals, Thailand, for providing blood samples. SKA also thanks CMU Presidential Scholarship for her financial support.

## Author contributions

SKA: conceptualization, formal analysis, investigation, methodology, visualization, writing—original draft, and writing – review & editing. SS: investigation, validation, review & editing. KD: investigation, review & editing. CP: conceptualization, funding acquisition, methodology, supervision, visualization, writing— original draft, writing – review & editing. All authors contributed to the article and approved the submitted version.

## Data and materials availability

All data are available in the main text or the supplementary materials.”

## Supplementary Materials

Materials and Methods

Figs. S1 to S8

Tables S1 to S7

