## Supplementary material for "Generation of Neural Stem/Progenitor-like Cells from Cultured Human Peripheral Blood Mononuclear Cells combined with Cannabidiol": Materials and Methods, Figs. S1 to S8

**The PDF file includes:**

Materials and Methods
Supplementary Text
Figs. S1 to S8
Tables S1 to S7

#### Materials and Methods

Peripheral blood mononuclear cells (PBMCs) isolation

PBMCs were isolated from peripheral blood of healthy adult volunteers aged 18-49 years old, of both sexes with Asian ethnicity and free from infectious disease (hepatitis B, hepatitis C, HIV) and no history of chronic disorders (inflammatory or autoimmune diseases). Informed consent was obtained from all participants. Ethical approval for PBMC experiments was obtained from the AMS Human Research Ethics Committee (AMSEC66EX008). Additionally, the buffy coat samples obtained from fresh blood donations at Blood Bank Unit of Chiang Mai University Hospital and Naresuan University Hospital (Thailand) were also used for PBMC isolation. PBMCs were isolated by density-gradient centrifugation using Lymphoprep™ (AXIS-SHIELD PoC AS) according to manufacturer's instructions. Red blood cells were removed using RBC lysis buffer (1x Ammonium chloride lysing buffer). The collected cells were suspended in RPMI1640 medium (Caisson Laboratories, USA) containing 10% fetal bovine serum (Gibco™), incubated at 37ºC in 5% $CO_2$.

Cell viability assay

PBMCs ($1x10^6$ cells in 200 µL) were treated with various doses of CBD. After 24, 48, and 72 hours of incubation, the cell suspension were then added with 20 µL of resazurin (0.1 mg/mL). Following a 4-hour incubation with resazurin, fluorescence emission intensity was measured using a spectrofluorometer at 590 nm (excitation wavelength: 570 nm).

Cell viability was calculated using the following equation:

$$\%\,Cell\ viability = \frac{F_{treat}}{F_{control}} \times\ 100$$

where $F_{treat}$ is fluorescence intensity in CBD-treated cells and $F_{control}$ is fluorescence intensity in untreated control cells.

Cell reprograming and differentiation in basic culture medium

PBMCs were seeded in RPMI1640 medium supplemented with various CBD doses (0 – 5 µg/mL), 10% heat-inactivated fetal bovine serum and 1% penicillin-streptomycin. The cultures were maintained in a humidified incubator set at 37ºC and 5% $CO_2$. Every seven days, a new medium was added. The cell morphology and differentiation behavior were examined under an inverted microscope (Nikon, Eclipse Ts2).

Immunofluorescence (IF) staining

Converting cells or differentiated cells were fixed in 4% paraformaldehyde for 10 minutes at room temperature. After fixing, the cells were permeabilized with 0.2-0.4% Triton X 100 (BioBasic, Canada) (if needed), and were blocked in 0.1% BSA (Capricorn Scientific, Germany) and in PBS at 37ºC for 15 minutes, followed by blocking with 2%-5% BSA at 37ºC for 1 hour. Primary antibody incubation was performed at 37ºC for 2 hours or 4ºC overnight. After washing with PBS, secondary antibody staining was performed at 37ºC for 60 minutes. Nuclei were counterstained with Hoechst33342 (APEXBIO, USA) and IF images were taken by fluorescence microscope (Nikon, Eclipse Ts2) and confocal microscope (Leica, Stellaris 5). The following antibodies were used to characterize and identify target proteins at different cell stages: Cell colony: OCT4 (Cat#653702, BioLegend), SOX2 (Cat#AB5603, Sigma-Aldrich),

TRA-1-81 (Cat#330710, BioLegend), PAX6 (sc-81649, Santa Cruz), Musashi-1 (Cat#AB5977, Sigma-Aldrich), Nestin (NES) (Cat#MAB5326, Millipore). Differentiation cells: SOX2 (Cat#AB5603, Sigma-Aldrich), PAX6 (sc-81649, Santa Cruz), Musashi-1(Cat#AB5977, Sigma-Aldrich), Vimentin (V9) (sc-6260, Santa Cruz), Nestin (NES) (Cat#MAB5326, Millipore), GFAP (Cat#HPA056030, Millipore), Synapsin I/II/II (Cat#853712, Biolegend), Tubulin ß3 (TUBB) (Cat#801208, Biolegend), NF-1 (sc-398751, Santa Cruz).

Alkaline Phosphatase assay

An Alkaline Phosphatase detection kit (Sigma-Aldrich) was used to check whether the colonies resemble stem cell colonies and the assay was followed the manufacturer's protocol. Briefly, the colonies were fixed with 4% paraformaldehyde in PBS for 1–2 minutes. After washing, the staining solution was added to the wells and incubated in the dark at room temperature for 15 minutes, then the number of red stained cell colonies was then observed under the inverted light microscope.

RNA sequencing

To check target gene expression and changes in lineage switching, developmental process, stem cells parameters and regenerative capacity, non-differentiated cells (freshly isolated) and the differentiated cells with and without CBD treatment were collected for RNA sequencing. The NucleoSpin RNA Plus kit (MACHEREY-NAGEL, Germany) was used to isolate total RNA from the cells. The quality and integrity of the total RNA was examined using agarose gel electrophoresis and an Agilent Bioanalyzer 2100 system.  Following a quality control check, mRNA purification and cDNA library construction were performed according to the manufacturer's instructions. Sequencing was conducted on the Illumina HiSeq PE150 platform. Next, the RNA-Seq data analysis was subsequently carried out. Quality control of raw data was performed using FastQC. Trimming was then done to the raw RNA-seq readings and STAR was used to map the clean reads to the hg19 human reference genome. The number of reads mapped to each gene was counted using the FeatureCounts program. Differential expression analysis between two samples based on DESeq2 was performed on a Nucleic Acid SeQuence Analysis Resource (NASQAR) with a DEseq2 Shiny tool. The differential expression genes (DEGs) were defined as the genes having a log2-transformed fold change of >2 between groups and an adjusted $P < 0.001$. Gene ontology (GO) enrichment analysis of the DEGs was also performed on NASQAR. In addition, the selected DEGs were analyzed for protein–protein interaction (PPI) networks using STRING software along with functional enrichment analysis.

Proteomic analysis

For cellular proteomic analysis, the untreated cells and CBD-treated cells (cultured in 6-well plates for 39 days (from 2 independent samples) were lysed in lysis buffer (1% SDS, 5mM dithiothreitol (DTT) in 20 mM HEPES-KOH (pH 8.0), 20 mM NaCl with 1x protease inhibitor) with incubation lasting 2-3 minutes. The lysed cells were transferred into 1.5 mL centrifugation tubes and were spun down at 12,000×g for 10 minutes. The supernatant was collected and kept at 4°C for a short waiting time. The protein concentration was adjusted to 2 mg/ mL with the same lysis buffer (volume 50 µL, total protein content = 100 µg). The tryptic peptides were analyzed and acquired using Orbitrap HF hybrid mass spectrometer combined with an EASY-nLC1000 nano-liquid chromatography (LC) system. Proteome Discoverer 2.4

was used to process the raw mass spectra (.raw file) and comparisons were made to the Uniprot protein database (organism: Homo sapiens). Protein identification and quantification were conducted using the following parameters: a minimum of three fragment ion matches per peptide; trypsin as the digest enzyme; cysteine carbamidomethylation as a fixed modification; and methionine oxidation as a variable modification. The peptide tolerance was set to 20 ppm; the fragment tolerance was set to 0.1 Da. The software's normalization technique (total intensity count) was used to normalize the ratio of relative protein abundances for each LC-run (across all runs; n = 10). The significantly enriched proteins in CBD-treated cells (padj <0.05), as compared to untreated control cells, were selected to perform functional enrichment analysis on the Database for Annotation, Visualization, and Integrated Discovery (DAVID). Different enrichment terms were categorized into different annotation clusters and enrichment scores were determined.

For secreted protein identification analysis, conditioned media collected from untreated cells and CBD-treated cells (cultured for 39 days) were lysed in lysis buffer (8 M urea in 100 mM Triethylammonium bicarbonate, TEAB). Proteins were reduced by 10 mM dithiothreitol (DTT) for 30 minutes at 37ºC and alkylated with 40 mM iodoacetamide (IAA). The protein lysates were digested with trypsin for 16 hours at 37ºC. The tryptic peptides were desalted with Pierce TM Peptide Desalting Spin Column and then analyzed with LC-MS/MS analysis. LC-MS/MS analysis was performed using an EASYnLC1000 system coupled to a Q-Exactive Orbitrap Plus mass spectrometer (Thermo Scientific, San Jose, CA). The separation was performed for 90 minutes at the flow rate of 300 nL/minute using EASY-SprayTM C18 column (Thermo Fisher Scientific). The stepwise gradient started at 5% solvent B and 95% solvent A, then proceeded as follows: 10% B for 20 minutes, 20% B for 40 minutes, and 40% B over the next 20 minutes. The gradient was finally held at 98% B for 2–8 minutes. Solvent A was 0.1% formic acid in $H_2O$, and solvent B was 0.1% formic acid in acetonitrile. The MS methods included a fullMSscan in the mass range of 350 – 1400 m/z at a resolution of 70,000, followed by 10 data-dependent MS2 scans at a resolution of 17,500. Mass spectra were searched against Human UniProt database (2022) using Proteome Discoverer 2.1 connected to SEQUEST-HT engine. Mass tolerance for precursor ions was set to 10 ppm and mass tolerance for fragment ions was 0.02 Dalton. Trypsin was specified with a maximum of 2 missed cleavages. Carbamidomethylation was chosen as a fixed modification, while oxidation of methionine was allowed as variable modification. A decoy search was performed using the false discovery rate of 0.01 as a threshold. Semiquantitative analysis of identified proteins was determined by measuring fragment area.

CBR expression by flow cytometry

To check whether the collected cells expressed ECS marker, CB2 expression was determined by flow cytometry. In a typical experiment, the cells fixed with 4% formalin for 15 minutes. After washing, the cells were blocked with 0.1% bovine serum albumin (BSA) for 30 minutes. After that, the cells were washed and then stained with primary antibody of cannabinoid receptor 2 (CNR2, Affinity Biosciences) at 4ºC overnight. After that, the cells were washed with PBS buffer and then incubated with Dylight 488 conjugated anti-rabbit secondary antibody (Invitrogen) at room temperature for 2 hours. After washing, the cells were subjected to analyse by flow cytometer (CytoFLEX, Beckman Coulter, USA).

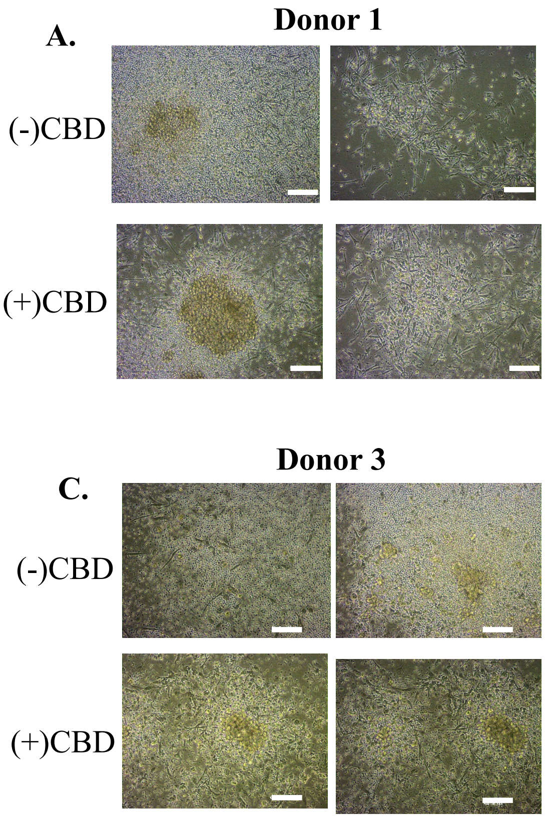


**Fig. S1.**

**Microscopic images (with two different areas) of PBMCs from four different donors** (A-D)**, cultured in 6-well plates for 7 days with and without CBD treatment. (Scale bar = 100 μm)**

### ECS vs Fetal NSPC

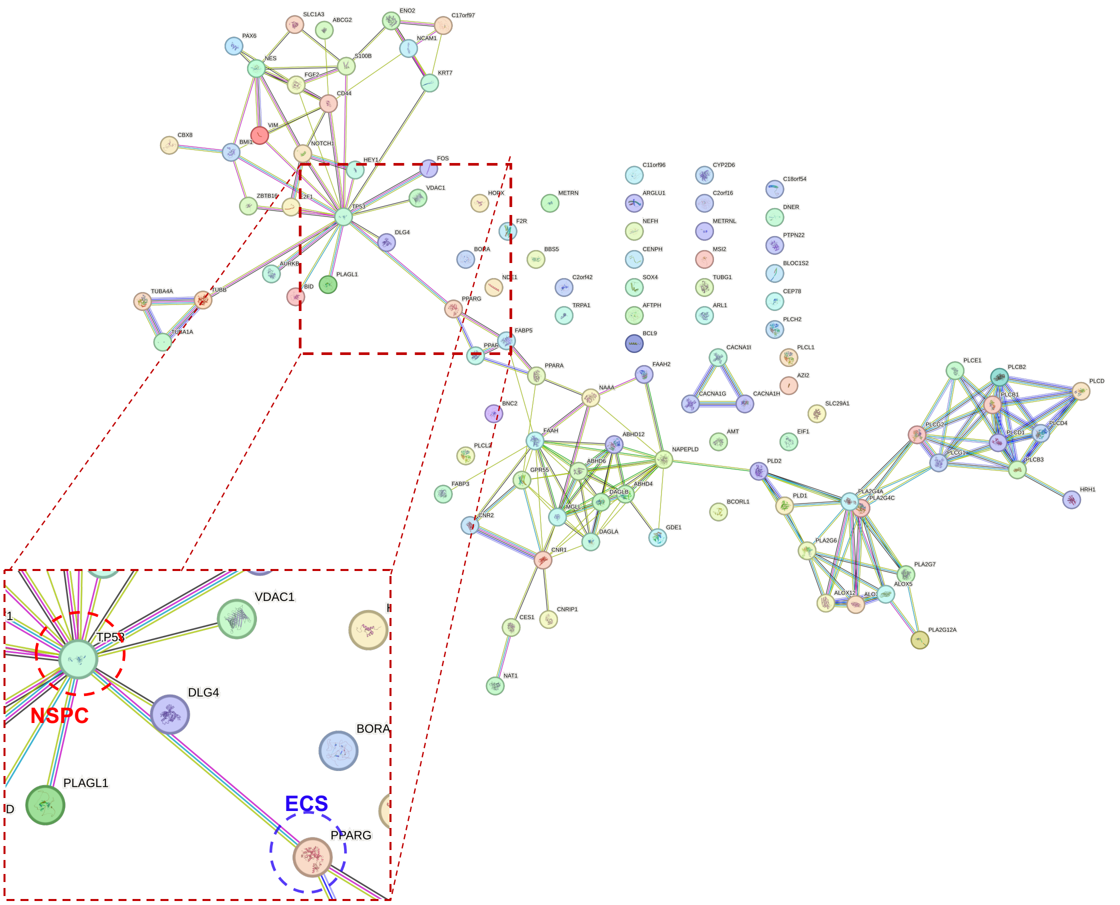


**Fig. S2.**

**PPI network (high confidence - 0.7), of the gene lists of ECS vs NSPC.**

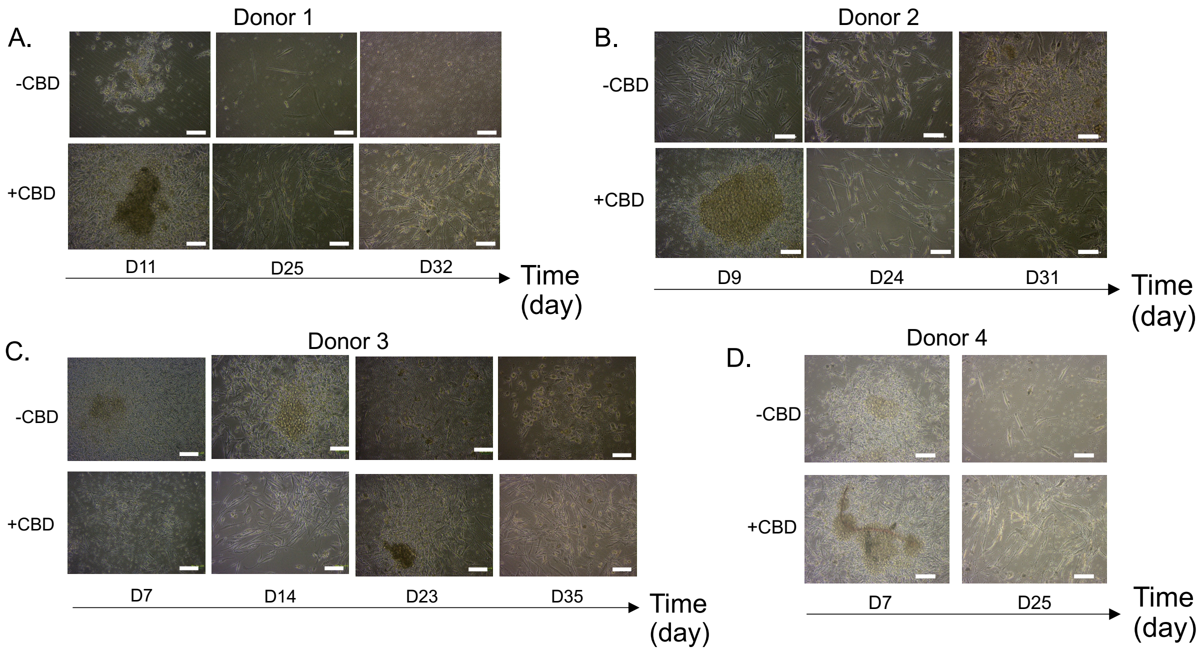


**Fig. S3.**

**Microscopic images of PBMCs from 4 different donors (A-D), cultured in 6-well plates for different lengths of time with and without CBD treatment. (Scale bar = 100 µm)**

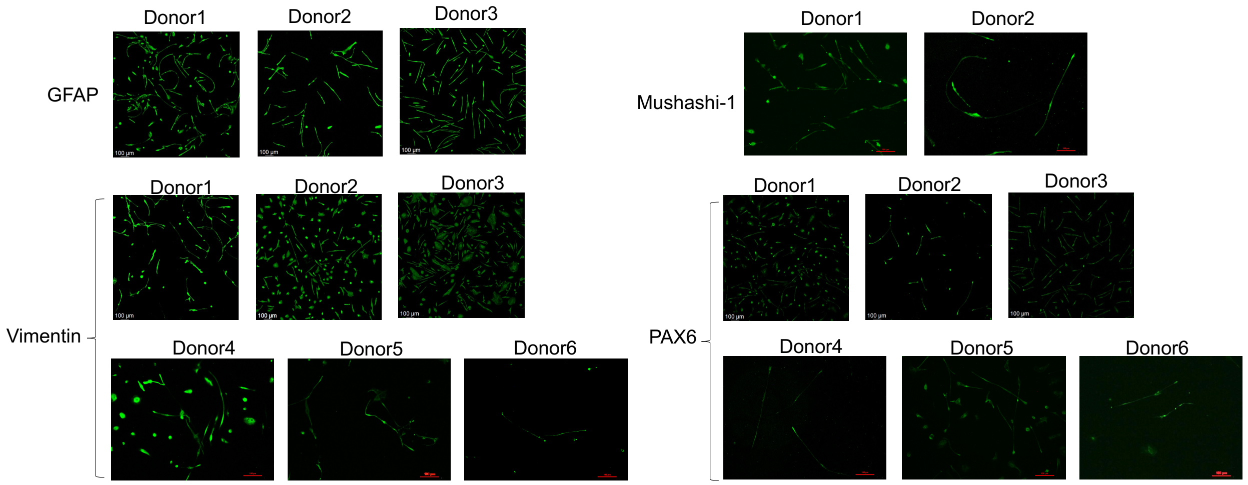


**Fig. S4.**

**Microscope images of immunofluorescent staining for NSPC markers (Vimentin, GFAP, PAX6, and Musashi-1) in differentiated cells (thin and long spindle shape cells) obtained from CBD treated PBMCs between different donors.**

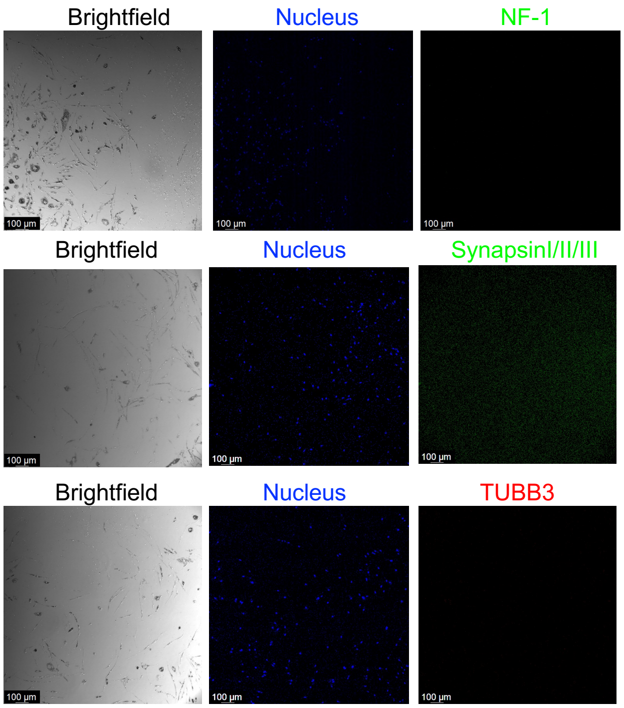


**Fig. S5.**

**Microscopic images of immunofluorescent staining for astrocyte marker (NF-1), neuron markers (Synapsin, TUBB) in differentiated cells (thin and long spindle shape cells) obtained from CBD treated PBMCs.**

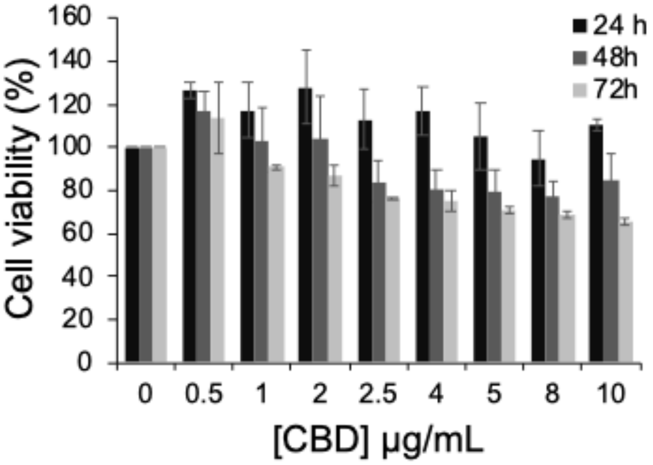


**Fig. S6.**

**Percentage of cell viability after treatment to various concentrations of CBD for different lengths of time.**

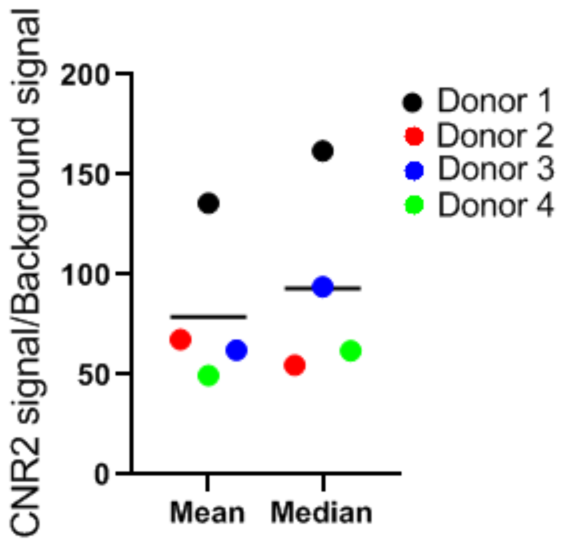


**Fig. S7.**

**Expression of cannabinoid receptor 2 on cell surfaces of PBMC from different donors.**

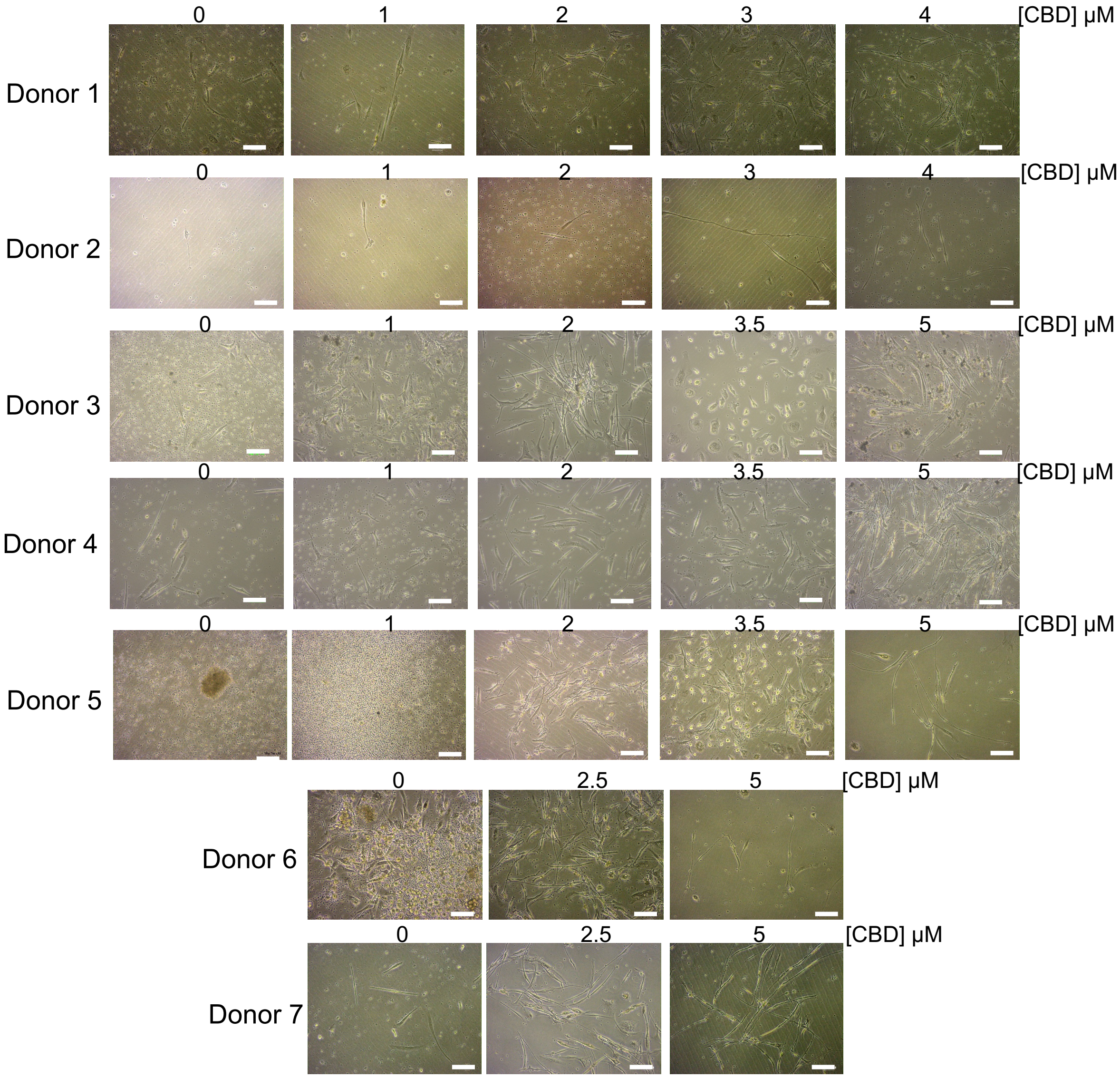


**Fig. S8.**

**Microscopic images of differentiated cells obtained from PBMC treated with CBD over 20 days at different doses of CBD between cells isolated from different donors. (Scale bar = 100 μm)**

**Table S1. Fetal NSPC gene set**

Tab 1A: Table of enriched DEG of fetal NSPC genes
Tab 1B: NSPC gene reference

**Table S2. Enriched fetal NSPC gene set**

Tab 2A: Table of enriched DEG of adult NSC genes
Tab 2B: Adult NSC gene reference

**Table S3. Enriched ECS DEGs of CBD treated cell**

Tab 3A: Table of enriched ECS DEGs of CBD treated vs untreated cell
Tab 3B: Reference gene list for ECS

**Table S4. List of identified proteins from proteomic analysis of CBD treated cells vs untreated cells**

**Table S5. Regenerative DEGs of CBD treated cells vs untreated cells**

Tab 5A: Table of enriched regenerative DEGs of CBD treated cells vs untreated cells
Tab 5B: Reference gene list for regenerative parameter

**Table S6. Enriched GO terms related to development, differentiation and stem cell of CBD treated vs untreated cell**

Tab 6A: Table of development related terms
Tab 6B: Table of differentiation related terms
Tab 6C: Table of stem cell related terms

**Table S7. List of plasticity and pluripotency DEGs in PBMCs without CBD treatment**
